# Relating layer fMRI signals to acoustics and intracranial neuronal activity in the human auditory cortex in a naturalistic design

**DOI:** 10.64898/2026.04.15.718614

**Authors:** Hsin-Ju Lee, Jyrki Ahveninen, Hsiang-Yu Yu, Chien-Chen Chou, Cheng-Chia Lee, Wen-Jui Kuo, Hankyeol Lee, Kamil Uludag, Fa-Hsuan Lin

## Abstract

How neuronal activity across cortical depths relates to fMRI signals during naturalistic perception has been difficult to verify noninvasively in humans. Here, we examine depth-dependent coupling between neuronal activity and fMRI signals during passive music listening. Depth-specific fMRI responses were modeled using envelopes of neural power in different frequencies elicited by the same naturalistic stimuli from a separate group of patients under intracranial EEG monitoring. From deep toward superficial depths, the relationship between neural power in different frequencies and fMRI signals systematically changed: alpha/beta activity (8-30 Hz) was increasingly associated with negative fMRI signals, while gamma-band (>30 Hz) activity showed increasingly positive associations. Gamma-band activity, including broadband high-frequency activity (>70 Hz), a proxy for neuronal firing, predicted fMRI signals beyond acoustic features, with coupling peaking at mid-to-superficial depths. Our findings reveal a systematic, depth-dependent pattern of neurovascular coupling in the human auditory cortex — a noninvasive link between neuronal frequency content and fMRI signals not previously shown in humans.

## Introduction

Anatomical connections of the human cortex differ across cortical depths to support feedforward or feedback information processing between cortical and subcortical regions [1]. Neurophysiological evidence from invasive recordings suggests that the feedforward and feedback processing are differentially manifested in neuronal oscillations [2–5] and broadband high frequency (>80 Hz) neural activities (HFA) [6,7]. Specifically, ascending feedforward pathways predominantly target intermediate cortical depths in primary sensory regions, while feedback pathways predominantly target superficial and deep depths [8,9]; correspondingly, gamma-band and high-frequency neural activity have been associated with feedforward or prediction-error signaling, while lower-frequency (alpha/beta) activity has been associated with descending feedback[2,10], consistent with the dominant oscillatory frequency changing across cortical depths reported by prior work [11–15].

This challenge can be mitigated by magnetic resonance imaging (MRI) for its non-invasive nature and submillimeter resolution. In particular, the intrinsic hemodynamic contrast in functional MRI (fMRI) is closely related to local field potentials [16–18]. Previous fMRI studies in the visual and sensorimotor cortices reveal that fMRI signals are negatively correlated with alpha and beta band oscillation power and positively correlated with gamma oscillation power [17,19–21]. During movie watching, the fMRI signal in the human auditory cortex was found to be positively and negatively correlated to gamma and alpha band neuronal oscillations, respectively [22]. However, because this previous intracranial work utilized standard-resolution fMRI, it could not resolve how these dynamics are organized across cortical depths. Conversely, recent advances using high-resolution 7T fMRI have elucidated depth-specific neurovascular coupling in the visual cortex using simultaneous scalp EEG [23]. Yet, scalp-EEG suffers from volume conduction and signal attenuation, severely limiting its ability to resolve the highly localized HFA. Consequently, how precise intracranial neural dynamics and vascular responses are coupled across cortical depths in the human auditory cortex remains elusive. Here, we bridge this critical gap by combining the high spatial resolution of 7T fMRI (0.8 mm isotropic) with the spectral precision of stereoelectroencephalography (SEEG) during naturalistic music perception.

Previous animal studies have revealed that neuronal activation in the primary auditory cortex differs across cortical depths in response latency [24,25] and selectivity to acoustic stimulus frequency (tonotopic organization) [25,26]. The representation of auditory information along the vertical axis of the auditory cortex also differs across cortical layers: the granular layers have a more accurate tonotopic response than supragranular and infragranular layers [27–29]. A recent high-resolution fMRI study in the human auditory cortex suggest that the preferred acoustic stimulus frequency is highly constant across cortical depths, while task demands can sharpen the acoustic-frequency tuning at superficial depth more than at intermediate and deep depths [30]. However, unlike this evidence on fMRI tuning to acoustic stimulus frequency, how depth-specific fMRI signals relate to the frequency content of the underlying neural activity in the human auditory cortex remains less clear.

Here, we use the neural power in different frequencies, derived from intracranial neuronal recordings in participants under presurgical monitoring, to predict depth-specific hemodynamic responses to the same naturalistic stimulus in an independent dataset acquired using 7T fMRI [23,31,32] in healthy participants. We aim to characterize how depth-specific neurovascular coupling in the human auditory cortex is associated with frequency-band neural activity during naturalistic music perception. Given the preferential anatomical targeting of intermediate cortical depths by feedforward connections in primary sensory regions [8,33], we predict that fMRI-neural coupling will be strongest at intermediate depths for frequency bands associated with feedforward signaling, and will be strongest at superficial and deep depths for frequency bands associated with feedback signaling. Specifically, based on prior work associating alpha- and beta-band activity with feedback modulation and adaptation at higher stages of the cortical hierarchy, and activity above 30 Hz (encompassing both oscillatory and non-oscillatory/broadband components) with feedforward processing of sensory input [34], we predict that fMRI coupling with alpha/beta-band activity and with activity above 30 Hz will show distinct depth-dependent profiles. Because feedback modulation is thought to become more prominent at higher stages of the cortical hierarchy, we additionally test whether these depth-dependent associations differ between the primary auditory cortex (A1) and its neighboring secondary auditory cortex (A2), which occupies a higher position in the auditory hierarchy.

## Results

**Figure 1A** shows the locations of implanted contacts around the auditory cortex. In this study, we included measurements on all contacts within 15 mm of the centroid of the left and right auditory cortex defined by an atlas derived from the Human Connectome Project [35]. This range was suggested by our previous SEEG study, which showed the correlation among electrode contacts separated within 20 mm [36]. Together, six and seven patients had contacts within 15 mm to the centroid of left (n=33) and right (n=29) auditory cortex, respectively (**Table S1**). Three (2 patients) and three (2 patients) contacts were within 5 mm to the centroid left and right auditory cortices, respectively. None of the electrode contacts were close to the epileptic spike generators, as confirmed by the neurologists. The atlas also provided the delineation of primary (A1) and secondary (A2) auditory cortices (**Figure 1B**). Because normalized depth (n.d.) is defined computationally via equidistant surfaces, these measurements represent geometric coordinates rather than absolute cytoarchitectonic boundaries. For clarity, we report our functional findings across ‘depths’, utilizing the BigBrain histological projection (**Figure 1C**) to provide qualitative anatomical context. Except n.d.=0.1, each normalized depth consisted of two to three cortical layers in the auditory cortex, A1, and A2. The depth n.d. = 0.5 included the largest proportion for layer IV: 36%, 61%, and 30% for the auditory cortex, A1, and A2, respectively.

**Figure 1.**
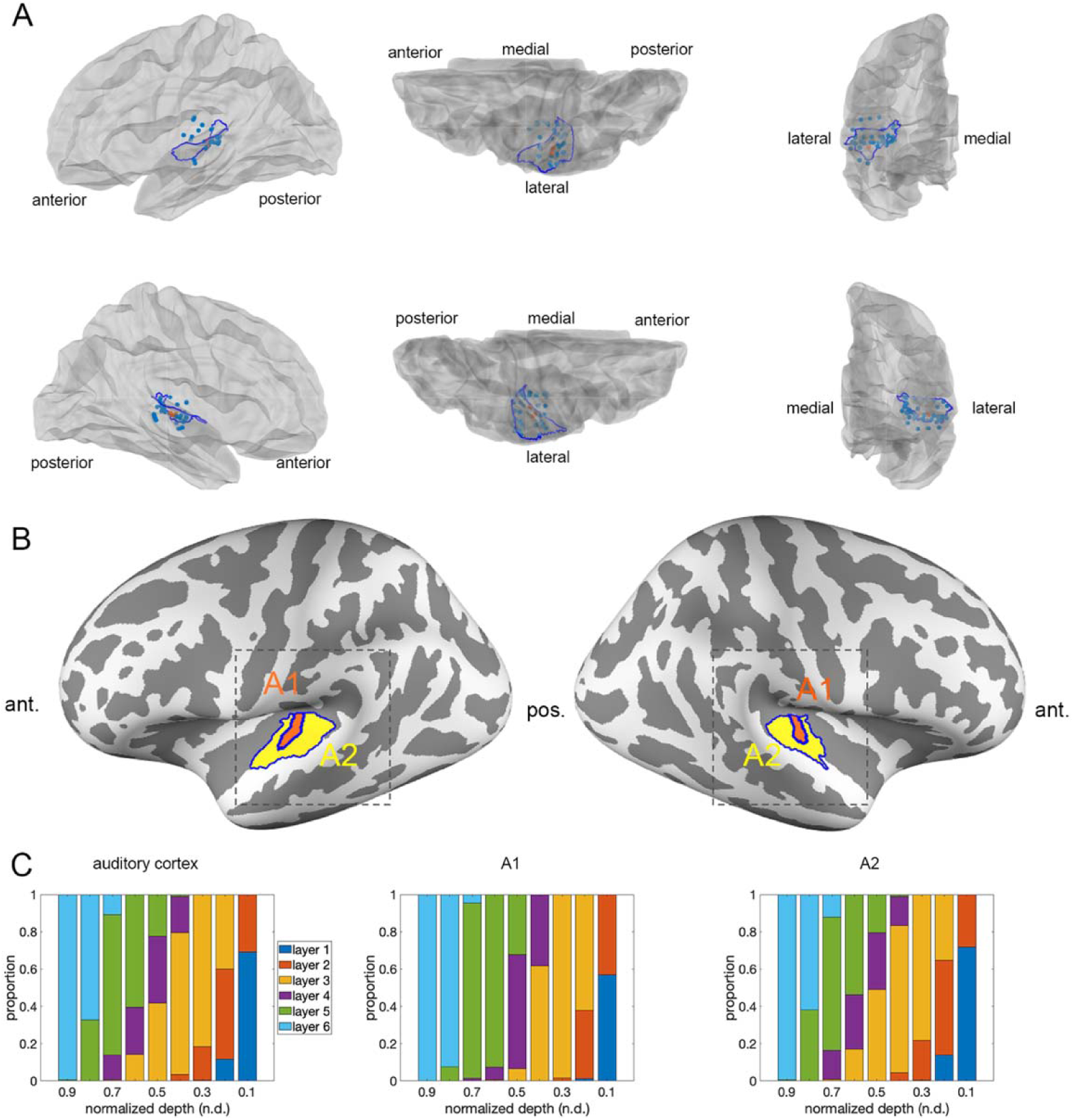
Implanted electrodes in the auditory cortex and the primary (A1) and secondary (A2) auditory cortex. (**A**) Blue dots denote the implanted electrode contacts from all epilepsy patients within 15 mm to the centroid of the auditory cortex (blue boundary) at the left and right hemispheres. Orange dots are electrode contacts within 5 mm to the centroid of the auditory cortex. (**B)** Boundary of A1 and A2 from the atlas. Ant.: anterior. Pos.: posterior. (**C)** Proportions of six cortical layers identified in Big Brain in 10 equi-distance boundaries between pial and gray/white matter in the auditory cortex, A1, and A2.

### fMRI signals entrained by musical stimuli vary across cortical depths

Following the experiment design of a previous study [22], our SEEG-fMRI correlation study used a cross-participant design: SEEG was measured from patients and fMRI was obtained from healthy participants. We first studied how the fMRI signals in the auditory cortex are entrained by acoustical features. Specifically, the depth-specific fMRI signals were correlated with the modeled fMRI signal generated by convolving the envelope of the presented music and a hemodynamic response function (HRF) [37]. **Figure 2A** shows the t-statistics of the correlation between the acoustic envelope and fMRI signals; only voxels significant at FDR-corrected p<0.05 are displayed. **Figure 2B** shows the mean Fisher-z partial correlation — a scale-free transform of each subject’s regression t-statistic (see Methods) — between the acoustic envelope and fMRI signals at each cortical depth, separately for the whole auditory cortex, A1, and A2 (subject-level, n=10). In **Figure 2A**, the spatial extent of significant vertices was largest at n.d.=0.1, n.d.=0.3, and the pial surface. In **Figure 2B**, the same pattern was visible in the subject-level mean correlation magnitude: n.d. = 0.1, n.d. = 0.3, and pial showed the largest mean Fisher-z values in all three ROIs (nominal p<0.05, uncorrected), compared to the deep depths (n.d.=0.5 to 0.9 and the white matter boundary). However, none of the seven depths reached FDR-corrected significance at the subject level (n=10; q≥0.07 in all three ROIs), so this depth-dependent pattern should be read as a consistent nominal trend rather than an independently established, subject-level effect.

**Figure 2.**
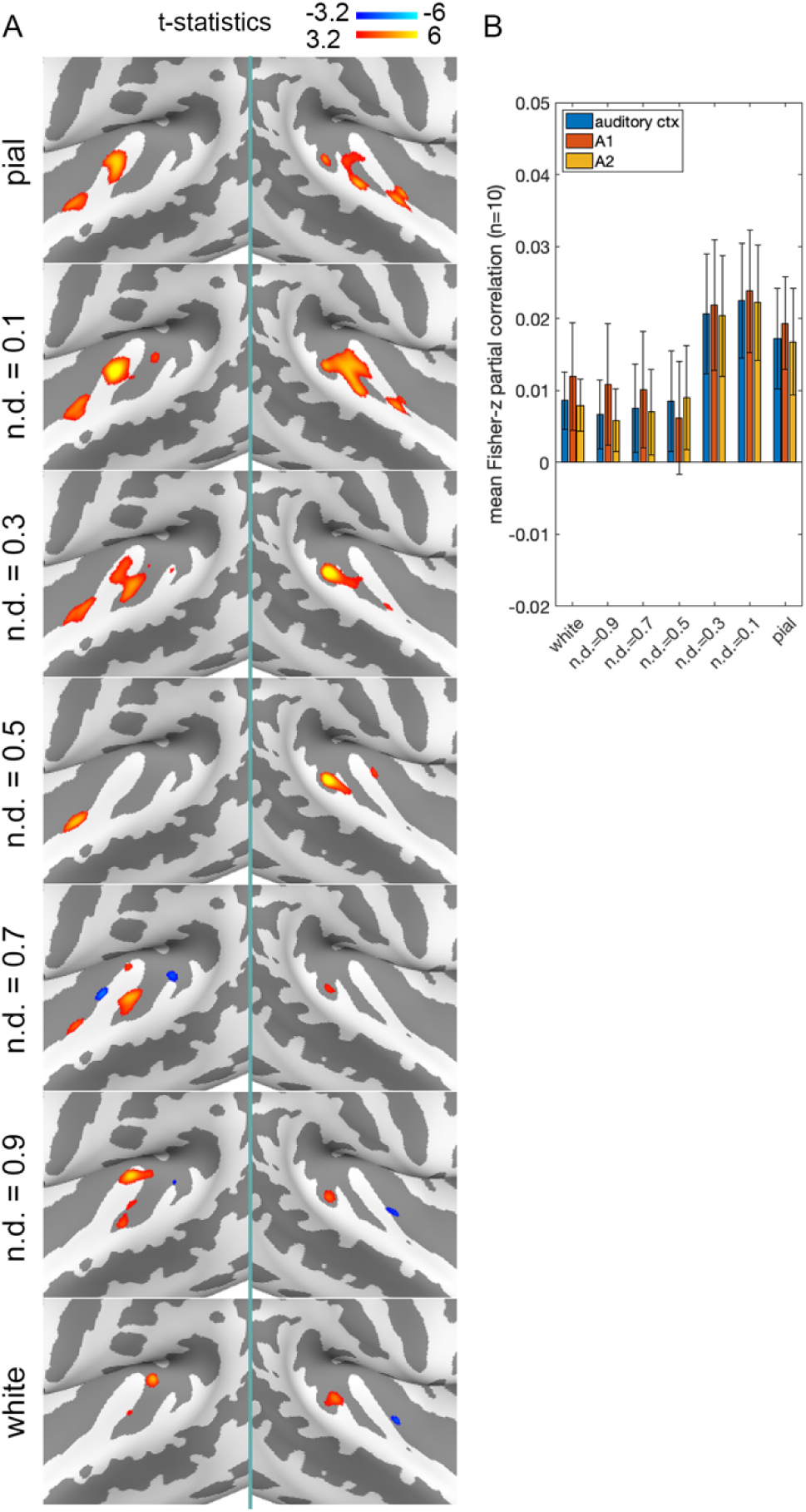
Correlations between music acoustical features and fMRI signals. (A) Distributions of the t-statistics of the correlation between the music acoustic envelope and fMRI signals across cortical depths (FDR-corrected p<0.05) around the auditory cortices in both hemispheres. See Figure 1 for the enlarged areas. (B) Mean Fisher-z partial correlation (subject-level, n=10) between the music acoustic envelope and fMRI signals at the auditory cortex (ctx), A1, and A2 across cortical depths. No depth reached FDR-corrected significance at the subject level, and a paired comparison between A1 and A2 found no significant difference at any depth, so no comparison brackets are shown.

At the same cortical depth, there was no significant difference in the acoustic-fMRI correlation between A1 and A2 (**Figure 2B**). We also compared the acoustics-fMRI correlation between cortical depths within each region, at the subject level (n=10, paired t-tests across all 21 unique depth pairs, FDR corrected). None of the 21 depth-pair comparisons reached significance in the whole auditory cortex, A1, or A2 (q≥0.07 throughout), despite a nominally significant trend at some depth pairs (uncorrected p<0.05) with moderate effect sizes (Cohen’s dz up to 1.26). We therefore do not report a specific pairwise-depth difference in the acoustics-fMRI correlation. The relationship between acoustic envelopes and fMRI signals across cortical depths was fitted to a cubic spline to create models for the auditory cortex, A1, and A2. These models included the coupling between stimulus acoustics and fMRI signal and potential bias caused by vascular signals.

### Correlations between fMRI and SEEG signals

We considered four canonical frequency bands: alpha (8–14 Hz), beta (14–30 Hz), low gamma (30–60 Hz), and high gamma (60–300 Hz). **Figure 3A** and **3B** show the between-subject [22] correlation between fMRI signals in the auditory cortex and the predicted responses generated by convolving band-limited SEEG activity patterns with a canonical HRF [37] during music hearing at different cortical depths. Significant negative correlations were found in alpha and beta bands at superficial (n.d.=0.1) and intermediate (n.d.=0.5) depths (FDR-corrected p<0.05). The envelope of gamma-band activity higher than 30 Hz convolved with hemodynamic response function (HRF) was positively correlated with fMRI signal in all depths. These depth-dependent associations should be interpreted cautiously: a sensitivity analysis correcting for potential deep-to-superficial venous signal leakage found that all four canonical frequency bands’ depth profiles, not only gamma, weaken and can reverse sign as the assumed strength of this correction increases, so no single depth profile can be treated as a fully correction-independent result (see Discussion). Between cortical depths, fMRI signals at the geometric midpoint (n.d.=0.5) were less negatively and less positively correlated with alpha/beta and gamma band activity than at the most superficial depth (n.d.= 0.1), respectively. Relative to the deep depth (n.d.=0.9), fMRI signals at the intermediate depth (n.d.=0.5) were more negatively and positively correlated with alpha and gamma band oscillations, respectively. This pattern across alpha, beta, and gamma bands is dominated by a single depth-dependent profile shared across frequency bands (a rank-one decomposition explains 98.9–99.7% of the squared coefficient magnitude in each region of interest), rather than indicating independent, band-specific neural generators localized to particular cortical depths. A dominant shared profile does not exclude a smaller frequency-dependent difference in depth-profile shape, which we tested directly (see below). The HRF-convolved band regressors were themselves correlated (alpha–high gamma r=−0.28, beta–high gamma r=−0.17, low–high gamma r=0.77), and the reconstructed depths were strongly collinear (variance-inflation factors up to ∼97), so no single depth carried a significant association with the SEEG signal once the other depths were modeled (smallest family-wise corrected p=0.48).

**Figure 3.**
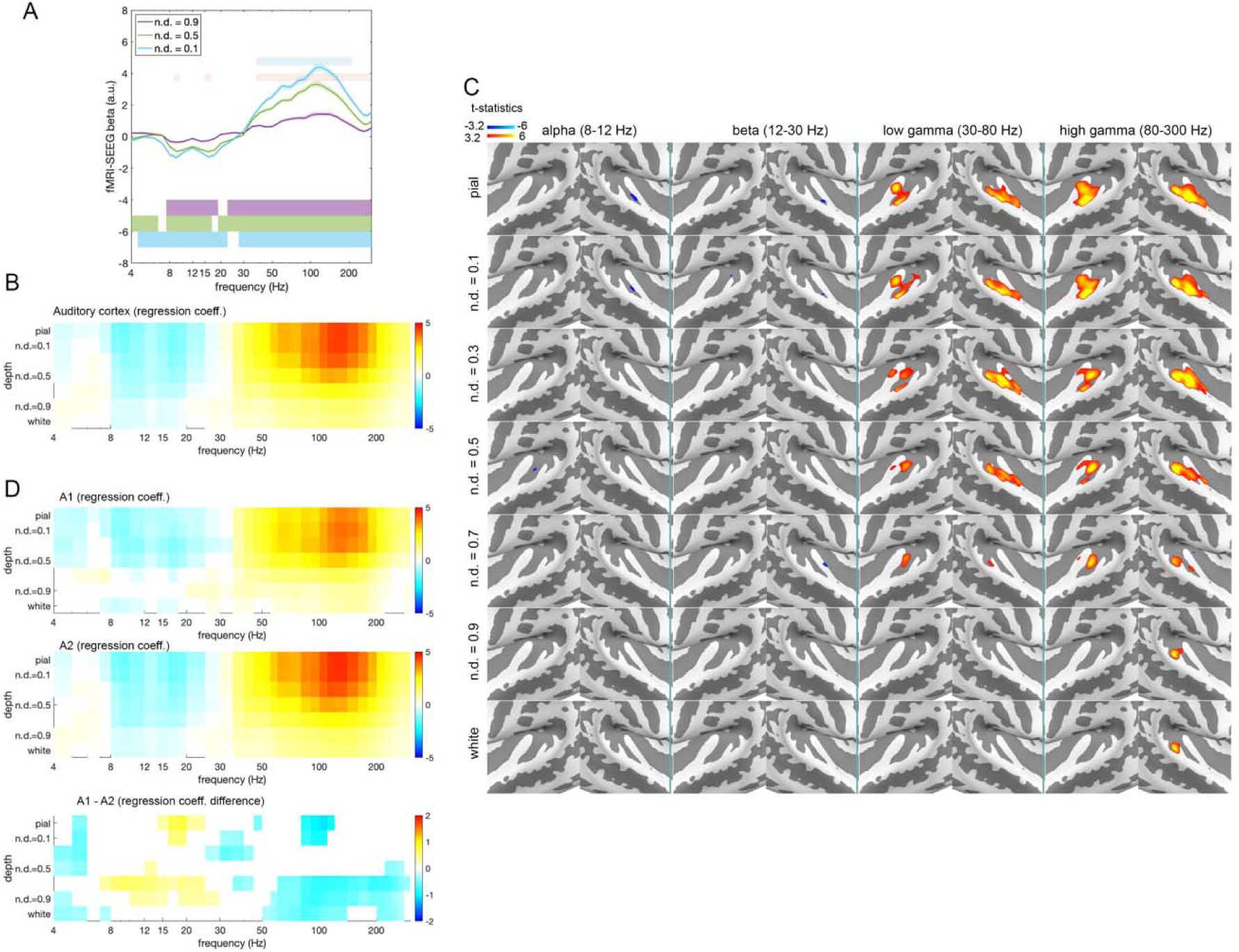
Correlations between intracranial neural activity and fMRI signals across cortical depths. Canonical frequency bands: alpha (8–14 Hz), beta (14–30 Hz), low gamma (30–60 Hz), high gamma (60–300 Hz). (A): The correlation between fMRI signals at three cortical depths (normalized depth, n.d., of 0.1, 0.5, and 0.9) and neural activity measured by electrodes within 15 mm to the centroid of the auditory cortex. Shaded bands: 95% bootstrap confidence intervals across vertices within the region of interest (1,000 resamples). Values are regression coefficients. Significances of regression coefficients (beta’s; false discovery rate, FDR, corrected p<0.05 across frequencies) at cortical depths are shown in horizontal bars at the bottom of the panel. The light blue and red horizontal bars at the top of the panel indicate significant differences between the three depths shown in this panel only (n.d.=0.5 vs. n.d.=0.9, and n.d.=0.5 vs. n.d.=0.1, respectively; FDR corrected p<0.05 across frequencies), not a comprehensive depth-pairwise comparison. these bars are from a descriptive, vertex-level test. (B): Regression coefficients between fMRI signals across cortical depths and neural activity. Significant coefficients (FDR corrected p<0.05 across frequencies and cortical depths) are color-coded. (C): Distributions of the t-statistics of correlations between intracranial neural activity and fMRI signals across cortical depths (FDR corrected p<0.05) in alpha, beta, low gamma, and broadband high frequency activity around the auditory cortices in both hemispheres. See Figure 1 for the enlarged areas. (D): Regression coefficients between neural activity and fMRI signals across cortical depths in the primary (A1) and secondary (A2) auditory cortices, and their difference (A1− A2). In the A1 and A2 panels, significant coefficients (FDR corrected p<0.05 across frequencies and cortical depths) are color-coded. In the A1 − A2 panel, colored cells mark where an unpaired two-sample t-test between A1 and A2 vertices was significant (FDR corrected p<0.05 across frequencies and cortical depths); because this vertex-level test treats spatially correlated vertices as independent, it is descriptive. A subject-level paired test (n=10) on scale-free Fisher-z partial correlations found a significant A1–A2 difference only in the high-gamma band at mid-to-deep depths (n.d.=0.5, 0.7, and 0.9; see Results).

### Frequency-dependent depth-profile shape

Because an omnibus rank-one test spreads its sensitivity over all possible departures from a shared profile, we also tested one specific difference chosen in advance, using the same depth-resolved coefficients (see Methods). A published drainage model [38] predicts the depth profile that activity in the middle cortical layers (III–IV) would produce; because blood drains toward the surface, this profile is weighted toward the upper half of the cortex. We asked whether two bands’ depth profiles differ from each other along this predicted pattern, after removing the profile they share. The test statistic is exactly zero whenever two bands share the same depth-profile shape, regardless of their amplitudes or signs; it therefore cannot be produced by a signal-to-noise difference that scales a band’s coupling uniformly across depths, nor by a venous drainage or spatial blurring operator common to all bands. Because the predicted pattern closely resembles the shared profile, this test is sensitive to a small residual component (about 5% of the pattern), which does not correspond to a simple superficial-versus-deep contrast. Alpha/beta- and gamma/HFA-coupled profiles differed along it in A1 (exact sign-flip test, p=0.002), but not in A2 (p=0.89) or across the whole auditory cortex (p=0.51). Bands sharing the same sign of coupling did not differ (A1: alpha versus beta, p=0.41; low gamma versus high gamma, p=0.86), although these pairs differ in signal-to-noise ratio by approximately 0.9 dB (SNR measured as band power during listening relative to each patient’s eyes-open resting recording, 9 patients; high gamma exceeded alpha by 2.61 dB, beta by 1.75 dB, and low gamma by 0.88 dB, Holm p≤0.047). The A1 result was unchanged when depth-dependent hemodynamic timing was modeled by adding the temporal derivative of each SEEG regressor (p=0.002). Smooth depth contrasts not derived from the drainage model, a linear gradient and a superficial-versus-deep step, did not detect a difference (p=0.58 and p=0.42). We therefore do not interpret the direction of this difference.

Distributions of significant correlations between depth-specific fMRI signals and neural activity in alpha, beta, and gamma bands are shown in **Figure 3C**. Much stronger and more diffuse patterns in the temporal lobes were observed in the gamma band than in the alpha and beta bands, with a trend of increasing correlations toward superficial depths. The correlations in alpha and beta bands are prominent in the left hemisphere.

The correlation between fMRI signals and SEEG signals in A1 and A2 was similar across cortical depths (**Figure 3D**). Both regions exhibited negative correlations in the alpha and beta bands and positive correlations in the gamma band. Using a scale-free measure of coupling strength (Fisher-z-transformed partial correlations, removing the effect of any difference in raw signal magnitude between A1 and A2) and a subject-level paired test (n=10, FDR corrected across the 7 depths and 4 canonical bands), A2 showed consistently larger-magnitude coupling than A1 across nearly every band and depth, but this difference reached significance only in the high-gamma band, at mid-to-deep depths (n.d.=0.5, 0.7, and 0.9); no other band or depth showed a reliable A1-versus-A2 magnitude difference at this sample size. In contrast, the depth-profile shape (independent of magnitude) was significantly correlated between A1 and A2 in the alpha, low-gamma, and high-gamma bands (mean shape correlation r=0.49–0.77 across subjects, FDR-corrected), indicating that A1 and A2 share a similar depth-dependent pattern even where their absolute coupling magnitude differs.

Because alpha-band activity has been reported to behave differently from beta and gamma bands in other paradigms [23,39], we examined alpha’s depth profile at the single-frequency level (8, 9, 10, 12, and 14 Hz) rather than only as one band-averaged profile. All five discrete alpha frequencies showed the same depth-profile shape — negative at every depth, becoming more negative from the white matter boundary toward superficial depths — with no evidence of the lower-alpha (8-10 Hz) versus upper-alpha (12-14 Hz) dissociation reported by [39]. We also tested whether each of the four canonical bands (alpha, beta, low gamma, high gamma) carries a unique association with depth-resolved fMRI signal beyond the other three, using a nested-model test (Holm-corrected across bands, at each depth, region, and condition). Only high-gamma activity showed a significant unique contribution — at all 7 depths in the auditory cortex and A2, and at 5 of 7 depths in A1 — while alpha, beta, and low gamma showed no significant unique contribution at any depth, in any region, or in any condition. These results indicate that, in this dataset, alpha’s association with depth-resolved fMRI signal is not separable from the combination of beta, low gamma, and high gamma, and do not support treating alpha as an independently informative, depth- or feedback-specific channel distinct from the other bands.

Neural activityderived from more electrodes was distributed over a wider range (15 mm from the auditory cortex centroid). Therefore, the specificity of the neural activity estimates may be compromised when improving the signal by averaging data from more electrodes. We studied this effect by limiting electrode contacts (*n* = 6; three in the left hemisphere and three in the right hemisphere) only within 5 mm of the auditory cortex. Repeating the correlation between fMRI and the more selective SEEG measurements shows similar results (**Supplementary Figure 1**). Namely, stronger negative correlations were found at the alpha and beta bands and positive correlations at the gamma band. Again, the correlations became more significant toward superficial depths. Distributions of the positively correlated gamma-band activity and fMRI signals were more focal when we limited to electrode contacts within 5 mm of the auditory cortex centroid. As with the main (15-mm) analysis above, this depth-dependent pattern should be interpreted subject to the same rank-one/shared-profile and venous-leakage-sensitivity caveats (see Discussion).

### HFA correlates fMRI signals across cortical depths

HFA taken from electrode contacts within 5 mm of A1 had significant correlation to depth-specific fMRI with a trend of stronger correlation toward the pial surface (**Figure 4A**). The strongest correlation between fMRI signals within A1 was at n.d.=0.3, while A2 and the whole auditory cortex had stronger correlation at n.d. = 0.3, 0.1, and pial surface (**Figure 4B**). Contrasting the HFA-fMRI correlation with the rescaled acoustic-envelope-fMRI correlation (**Figure 2**), a fixed-effect, vertex-level test — which describes a feature of this specific, already-collected dataset rather than testing whether the pattern generalizes to new participants — found that in the auditory cortex, HFA-fMRI correlation was significantly stronger than the acoustic-envelope-fMRI correlation at intermediate-to-deep depths (n.d.=0.5, 0.7, and 0.9) and significantly weaker at the superficial depth (n.d.=0.1); A2 showed the same pattern. In A1, HFA-fMRI correlation was significantly stronger at n.d.=0.5, but the corresponding differences at the white matter boundary and at n.d.=0.1 did not survive correction. A complementary, random-effect test at the subject level (n=10, paired t-test and exact sign-flip permutation) found no significant HFA-versus-acoustic-envelope difference at any depth or region after FDR correction (all p≥0.12); this design has 80% power only for effect sizes (Cohen’s dz) at or above 1.00, and every observed effect size falls well below that (range 0.01–0.54), so this null result should be read as an inability to determine whether the fixed-effect pattern generalizes to new participants, not as evidence that no difference exists. Unlike the corresponding acoustic-envelope depth-by-depth contrast (not retained; see above), the HFA-fMRI depth-by-depth contrast survives correction at the subject level (n=10, paired t-tests across all 21 unique depth pairs, BH-FDR corrected): 15 of 21 depth pairs were significant in the auditory cortex, 11 of 21 in A1, and 16 of 21 in A2. Coupling peaked at the mid-to-superficial depth (n.d.=0.3 in A1; n.d.=0.3 to the pial surface in the auditory cortex and A2), and the most significant depth-pair differences were between this peak and intermediate-to-deep depths (n.d.=0.7, 0.9, and the white matter boundary) (**Figure 4C**). The centroid of the area showing significant HFA-fMRI signal correlation was found moving from A1 toward A2, in the postero-lateral direction at deeper cortical depths (**Figure 4D)**.

**Figure 4.**
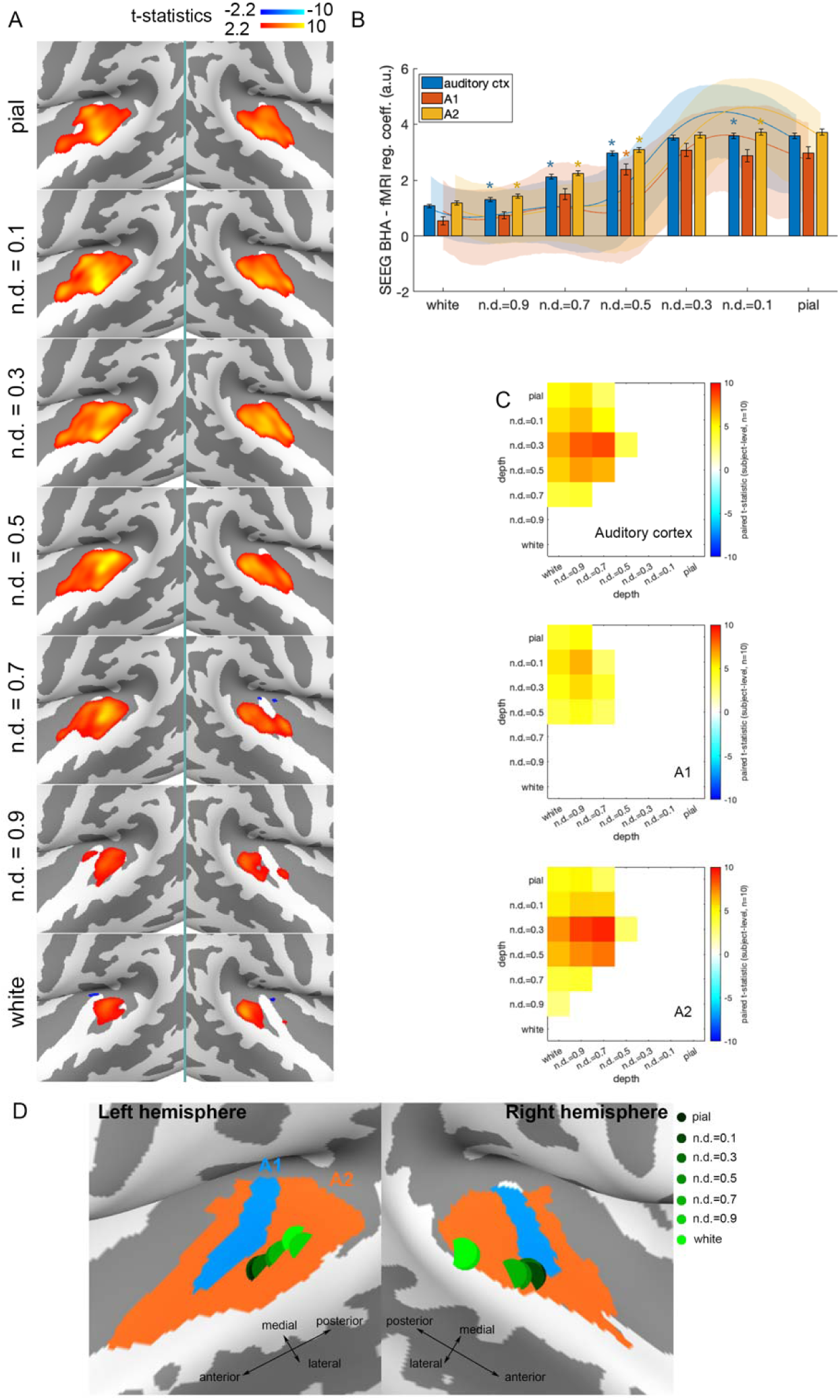
Correlations between HFA (70–150 Hz) and fMRI signals. (A): distributions of the t-statistics of correlations between HFA and fMRI signals across cortical depths (FDR corrected p<0.05) around the auditory cortices in both hemispheres. See Figure 1 for the enlarged areas. (B): Regression coefficients between HFA and fMRI signals at the auditory cortex (ctx), A1, and A2 across cortical depths. A paired comparison between A1 and A2 found no significant difference at any depth, so no comparison brackets are shown. Solid lines represent a cubic spline fit to the acoustic-envelope-fMRI correlation (Figure 2), rescaled to match the HFA profile’s amplitude, with a subject-level (n=10) bootstrap 95% confidence band (shaded). Colored asterisks (blue, orange, yellow) denote depths where a fixed-effect, vertex-level test (see Methods) found the HFA-fMRI correlation (bars) significantly different from the acoustic-envelope-fMRI correlation (lines) for the respective regions; this fixed-effect pattern was not confirmed by a complementary subject-level test (see Results). (C): The pair-wise, subject-level (n=10, paired t-test, BH-FDR corrected) comparison of HFA-fMRI correlation between cortical depths at the auditory cortex, A1, and A2. Values are t-statistics. Significant differences at data grids in the upper-left triangular part of the data matrix are color-coded. The value at row *r* and column *c* represents the difference (depth *r* − depth *c*). (D): The centroid of the area with significant correlation between HFA and depth-specific fMRI signals.

We also asked whether HFA-fMRI coupling is redundant with acoustic-envelope-fMRI coupling, using a nested-model comparison: after controlling for a rich acoustic-feature model (broadband envelope plus 8 spectral bands, 50–8,000 Hz), does adding the SEEG band regressor still significantly improve the fit to depth-resolved fMRI signal? Low-gamma and high-gamma activity showed a significant, largely depth-wide improvement in fit beyond the acoustic model, at nearly every depth in the auditory cortex and A2 (Holm-corrected across bands); alpha and beta did not show a comparably reliable unique contribution. This indicates that HFA/gamma-band SEEG activity is not redundant with acoustic-envelope-driven fMRI coupling — it carries information about depth-resolved fMRI signal that the acoustic-envelope model does not capture — complementing the depth-by-depth comparison above. Separately, we asked whether the shape of the HFA depth profile (its relative pattern across depths, independent of magnitude) is distinguishable from the shape produced by acoustic-frequency-driven fMRI coupling generally: comparing each canonical SEEG band’s depth-profile shape against 84 acoustic-spectrogram channels (4 Hz–8 kHz) processed through the identical pipeline, every band’s shape closely matched at least one acoustic channel (permutation p<0.05 in all 12 region-by-band tests). The matching acoustic frequency was neither consistent across regions nor close to the neural band’s own frequency, so this indicates that depth-profile shape throughout this dataset is dominated by a factor common to nearly any regressor passed through this pipeline (most plausibly the shared depth-dependent gain profile described above), rather than a feature distinctive to neural signals. This shape-genericity finding is compatible with, not contradicted by, the locally significant magnitude differences reported above: a generic depth-profile shape can still coincide with a specific, depth-resolved magnitude difference from one particular acoustic regressor, and with a significant unique contribution to fMRI variance beyond that regressor.

### Relationships of neural activity and fMRI signals across cortical depths change between repeated listening

As we measured SEEG and fMRI with repeated music presentation, we examined how the relationship between neural activity and fMRI signals across cortical depths changes between the first and second listening. We hypothesized that repeated listening might differentially affect this SEEG-fMRI relationship across cortical depths, frequency bands, and region, potentially reflecting contextual or predictive processing that could differ between A1 and A2 across the auditory hierarchy. **Supplementary Figure 2** shows the difference in SEEG-fMRI coupling between the first and second presentations, separately for A1 and A2 (vertex-level test, FDR corrected p<0.01 across frequencies and cortical depths). Visually, contrasting between the first and second listening, we found stronger negative correlations between alpha/beta band activity and fMRI signals and stronger positive correlations between neural activity above 70 Hz up to 200 Hz and fMRI signals in both A1 and A2. The significant differences were found more across frequencies, particularly between 50 Hz and 200 Hz, in A2 than A1. At A1, more significantly negative correlations in alpha and beta bands were found at intermediate (n.d.=0.5) depths. At A2, more significantly negative alpha/band-fMRI correlations were observed across cortical depths. Significantly stronger correlations at broadband high frequency activity (> 70 Hz) and fMRI signals were found across cortical depths with stronger effects above n.d.=0.5 at A1 selectively, while this difference was found across all depths at A2. Interestingly, we also found that the correlation between low gamma band (between 30 Hz and 50 Hz) activity and fMRI were reduced in the repeated listening, particularly around intermediate and superficial depths, at both A1 and A2. At the same time, a stronger correlation between theta band (between 4 Hz and 8 Hz) activity and fMRI signals was found at A1 and A2. This comparison does not contradict feedback-based models, but it cannot distinguish feedback-related neural adaptation from repetition-related changes in neurovascular coupling; moreover, because the SEEG and fMRI measurements come from different participant groups, it cannot establish within-person neural adaptation.

## Discussion

We studied the relationship between intracranial neuronal activity and depth-specific fMRI signals in the human auditory cortex during perception of complex naturalistic stimuli. Leveraging invasive recordings from epilepsy patients, we achieved high spatial resolution measurements of local field potentials. Combined with ultrahigh-resolution 7T fMRI (0.8 mm isotropic), this approach enabled us to characterize neurovascular coupling across primary and secondary auditory cortices during music processing.

The correlation between acoustic features and fMRI signals spanned all cortical depths, consistent with recent laminar recordings in the mouse auditory cortex [40]. The strongest absolute correlations between fMRI signals and the envelope of musical stimuli were found at mid-to-superficial depths (n.d.=0.3 and 0.1) (**Figure 2**), likely reflecting the general vascular draining bias toward the cortical surface. Because this macroscopic vascular bias obscures the underlying neural directionality, we leveraged our intracranial recordings to investigate how neural activity in distinct frequency bands relates to these depth-dependent hemodynamic responses.

In this frequency-band analysis, we found that fMRI signals correlate negatively with alpha/beta bands and positively with the gamma band, consistent with a correlational study of intracranial recordings and fMRI in the human auditory cortex during movie watching [22]. These correlations are more prominent at superficial depths than at the deep white-gray matter boundary (**Figure 3**). This depth-dependent pattern should be interpreted with caution: a sensitivity analysis correcting for potential deep-to-superficial venous signal leakage found that the pial-versus-white contrast for all four canonical frequency bands weakens and can reverse sign as the assumed strength of this correction increases, so we cannot identify a venous-leakage-correction-independent depth profile for any band (see Methods). The coupling between band-limited neural activity in alpha, beta, and gamma bands and fMRI signals did not show selective peaks at intermediate, superficial, or deep depths. Instead, a strong correlation (positive in gamma and negative in alpha/beta bands) was found at the superficial depth with a diminishing gradient toward deeper depths. These oppositely signed associations for alpha/beta versus gamma might appear consistent with predictive-coding frameworks that associate feedforward processing with gamma-band activity and feedback processing with alpha/beta-band activity [41]. However, the canonical microcircuit model underlying this framework [10] predicts that feedforward- and feedback-associated frequencies should be differentially distributed across cortical depth, not merely oppositely signed at every depth. A rank-one decomposition of the frequency-by-depth coupling matrix found that the four canonical bands are dominated by a single shared depth-dependent profile, with frequency-specific scaling and sign. A test targeted at a pattern derived from a published drainage model detected only a small departure from this shared profile, confined to A1 and not corresponding to a simple superficial-versus-deep contrast (see Results). This does not show the differential depth manifestation the predictive-coding framework would predict, so we do not interpret these results as support for that framework.

The neurophysiological basis of fMRI signal has been attributed to local field potentials in non-human primate studies [16,17]. These neural activities coupled to fMRI signals were represented in oscillations faster than 50 Hz [17,22]. Notably, HFA (>70 Hz) has traditionally been taken as a proxy for asynchronous multi-unit activity and local neuronal spiking [42,43]. However, recent evidence suggests that HFA is not a monolithic signal; it comprises distinct early components related to local spiking and late components driven by dendritic processes [7]. Our HFA results (**Figure 4**) highlight the complexity of this relationship across the cortical column. Our HFA-fMRI depth profile instead peaked at the mid-to-superficial depth (n.d.=0.3), differing most sharply from intermediate-to-deep depths (**Figure 4C**) — a location similar to where the purely acoustic envelope-fMRI correlation also peaks (**Figure 2**), consistent with the largely shared, non-frequency-specific depth-dependent gain profile described above rather than with a depth-segregated feedforward/feedback mechanism. Note that the link between synaptic input and fMRI signals was also supported by the anti-correlation in alpha and beta bands [2] (**Figure 3**).

Comparing A1 and A2 (**Figure 3D**), we found that A2 showed significantly larger-magnitude coupling than A1 specifically in the high-gamma band at mid-to-deep depths (n.d.=0.5, 0.7, and 0.9); no other frequency band or depth showed a reliable magnitude difference between these regions at this sample size (see Results). This more limited pattern does not support a general claim that feedback modulation is more prominent at higher stages of the auditory hierarchy across all frequency bands; it is specific to high-gamma-band coupling at these depths. The depth-profile shape, independent of magnitude, was similar between A1 and A2 in most bands (see Results), further indicating that these two regions are not clearly dissociated by a feedback-specific frequency signature in this dataset.

The neurovascular coupling studied here exhibited a general trend of a stronger correlation at superficial depths, even with our efforts of regressing out the average fMRI signal at pial and gray/white matter boundaries. This can be attributed to the venous flow from deep to superficial depths [44]. This feature has also been observed in EEG-fMRI correlations in the human visual cortex [23]. However, our analyses also suggest depth-specific properties: in the pairwise, subject-level comparison of fMRI-SEEG HFA correlation across depths (**Figure 4C**), the largest differences were found between the mid-to-superficial peak (n.d.=0.3) and intermediate-to-deep depths (n.d.=0.7, 0.9, and the white matter boundary). Furthermore, in the comparison between repeated listening (**Supplementary Figure 2**), venous bias was also reduced because this within-region vascular bias was not likely to change between conditions [45]. The reduced negative coupling between alpha/beta activity and fMRI signals may reflect a mechanism that prioritizes auditory processing [46]. The increased coupling between theta activity and fMRI signals may follow from reduced cognitive load during repeated listening [47]. A similar strategy of contrasting between conditions was used, taking the fMRI-acoustics correlation as a reference: the fMRI-SEEG HFA correlation differs from the fMRI-acoustics correlation mostly at intermediate-to-deep depths (n.d.=0.5, 0.7, and 0.9) and the superficial depth (n.d.=0.1) in a fixed-effect, vertex-level test (**Figure 4B**), although this pattern was not confirmed at the subject level (see Results). More complicated methods, such as biophysical modeling [48], may suppress the venous bias more specifically.

Depth-specific fMRI has been used most frequently in visual [49–51] and somatosensory cortices [52]. In auditory cortices, it has been shown that the tonotopic mapping remains stable across cortical depths, while attention can tune acoustic frequency preference at more superficial depths than at intermediate and deep depths [30]. Our results show how depth-specific fMRI signals are correlated with alpha, beta, and gamma band activity across the auditory cortex hierarchy. Together, we showcase how to use sub-millimeter fMRI to enrich the understanding of the neurophysiological underpinning of auditory perception and cognition.

A limitation of our approach is the translation of computationally defined normalized cortical depths to true cytoarchitecturally-defined cortical layers. While we qualitatively matched our equidistant surfaces to the BigBrain dataset [53] —showing, for instance, that layer IV is most heavily represented at n.d.=0.5 —cortical thickness and curvature variations mean that a depth like n.d. = 0.3 captures a blend of supragranular and granular features depending on the specific location (**Figure 1C**). Future quantitative studies using subject-specific high-resolution quantitative MRI are needed to achieve exact histological correspondence.

This study used musical stimuli, a potent acoustic stimulus known to influence emotions through both the auditory and limbic systems [54], to investigate how neuronal activity and blood flow are linked across different depths of the auditory cortex. This design echoes the increasing interest in exploring human brain function with more ecologically relevant stimuli [55]. Our stimuli included instrumental and vocal components in different genres, leading to the cortical-depth specific correlations that are expected to be generalized to other musical stimuli.

Our study used SEEG and fMRI data from different participants. This between-subject design [22] carries the advantage of avoiding the carryover effect (between SEEG and fMRI) but cannot control the tentative confound attributed to the difference between healthy participants providing fMRI measurements and epilepsy patients providing SEEG recordings. Taking SEEG and fMRI from different individuals was due to the practical challenges of scheduling epilepsy patients for 7T MRI. Note that to reduce the concern of effect from groups, all SEEG data from contacts showing epileptic features were excluded by clinicians. Future studies with SEEG and fMRI on the same patient can further address this concern.

We also did not perform a baseline-or task-contrast analysis (e.g., music versus a silent baseline) to more directly test feedforward and feedback processing. The only candidate baseline in this design is the approximately 10-second silent interval embedded before each song, which is too brief relative to the hemodynamic response to isolate a clean pre/post-stimulus contrast without substantial carryover from the preceding music. A properly controlled feedforward/feedback contrast would require a dedicated silence or low-level control condition of adequate duration, which was not part of this study’s design; we view this as a direction for future work rather than an analysis the present dataset can support. Relatedly, because a resting-state fMRI recording matched to the SEEG sessions is not available, we did not test whether the SEEG-fMRI coupling reported here differs between music listening and a resting baseline; our analysis characterizes coupling during music listening only, not a music-versus-rest contrast.

Taken together, our findings demonstrate that depth-specific fMRI, when combined with intracranial electrophysiology, can reveal a systematic, depth-dependent pattern of neurovascular coupling in the human auditory cortex during naturalistic perception. By relating frequency-band neural activity to hemodynamic responses across cortical depths, we provide a noninvasive window onto neurovascular coupling in humans, which we relate to, but do not use to establish, the feedforward and feedback processing streams described in prior work. This approach not only bridges invasive and non-invasive modalities but also opens new avenues for investigating the neural basis of complex cognitive functions and their disruption in neuropsychiatric disorders.

## Materials and Methods

### Participants

All epilepsy patients and healthy participants joined this study with written informed consent after the approval of the Institutional Review Board from Taipei Veterans General Hospital and SungKyunKwan University, respectively. All participants had normal hearing.

#### Healthy participants (fMRI)

Ten healthy participants underwent 7T fMRI while listening to the musical stimuli described below.

#### Epilepsy patients (SEEG)

Electrophysiological responses were measured from 16 medically refractory epilepsy patients by SEEG. Locations of the implanted electrodes were planned based on each patient’s clinical need. All electrode contacts showing epileptic-like activity, verified by neurologists, were excluded from the analysis. We excluded data from three patients because they had no contacts within 15 mm of the centroid of the left or right auditory cortex, defined from an atlas [35]. SEEG data were therefore analyzed from 13 patients and 62 electrode contacts for the primary (15-mm) selection; a more spatially restricted 5-mm selection (6 contacts, 4 patients) was also analyzed (see the a priori rationale under Statistical Analysis, below).

### Acquisition Protocols

#### Stimulus presentation

We presented three songs (song 1: Japanese animation Doraemon theme, 176 s; song 2: Brahms Piano Concerto No. 1, 360 s; song 3: Lost Stars from Adam Levine, 266 s) to each individual during fMRI or SEEG recordings, controlled by Psychtoolbox in MATLAB. These musical stimuli are available at https://github.com/fahsuanlin/labmanual/wiki/32:-Sample-data:-SEEG-recording-during-music-listening. Each participant was instructed to relax and listen to the musical pieces, each played twice in a randomized order. A 10-s silence interval was inserted as a rest condition between two songs, before the first song, and after the last song. Songs were played through the same MRI-compatible earphone (Model S14, Sensimetrics, Gloucester, MA, USA) in both modalities. In the fMRI session, music onset was synchronized to the first slice excitation of the first acquired volume. In the SEEG session, music onset was controlled by Psychtoolbox 3.0, which played the audio and simultaneously sent a trigger to the SEEG recorder at stimulus onset; the played audio was additionally fed back into an auxiliary SEEG recording channel, giving a continuous, sample-level record of the actual acoustic waveform, used post hoc to verify and refine alignment between the presented song and the SEEG recording.

#### fMRI acquisition

For healthy participants, functional MRI data were acquired on 7T MRI (Terra, Siemens Healthineers, Erlangen, Germany) with a 32-channel whole-head coil array (Nova Medical, Wilmington, MA, USA). Structural and functional images, both with 0.8 mm isotropic resolution, were acquired with MP2RAGE [56] and gradient-echo EPI (TR=2.8 s, TE=29 ms, flip angle=75°, 64 slices), respectively. The fMRI covered the temporal lobe at both hemispheres. The fMRI acquisition was “blip-down,” where the phase-encoding direction was from the positive to the negative direction. For each participant, we additionally acquired EPI “blip-up” images, with phase encoding from negative to positive direction. The combination of “blip-down” and “blip-up” trajectories allowed estimation of the field map, used to correct image geometric distortion (see Preprocessing, below).

#### SEEG acquisition

All patients had an electrode (Ad-Tech Medical Instrument, Oak Creek, WI, USA) with up to ten contacts (5 mm separation) implanted in the temporal lobe. Pre-surgery and post-surgery structural MRIs (3D T1-weighted images with MPRAGE sequence; 3T MRI, Skyra, Siemens Healthineers, Erlangen, Germany; 1 mm isotropic resolution) were obtained from all patients to identify electrode and contact locations (see Preprocessing, below).

### Preprocessing

#### fMRI (healthy participants)

Structural images were reconstructed to delineate the gray-white matter boundary via automatic segmentation and cortical parcellation using FreeSurfer (http://surfer.nmr.mgh.harvard.edu; version 7.2) . Nine cortical surfaces with equally spaced cortical thickness were reconstructed (*mris_expand* in FreeSurfer). In this study, we used “normalized depth” (n.d.) to denote the interpolated surface. For example, n.d.=0.1 was the surface about 10% from the pial surface and about 90% from the gray-white matter boundary. The relationship between normalized depths and cortical layers was quantified by morphing cortical layer thickness [53] from the BigBrain data, a 3D histological model of the human brain with 20-µm resolution [57], to the FreeSurfer atlas using the BigBrainWarp toolbox [58] (https://github.com/caseypaquola/BigBrainWarp). While the equi-volume principle is theoretically advantageous for accounting for cortical folding in cytoarchitecture, we opted for the equi-distance approach (*mris_expand*): at our in-vivo spatial resolution (0.8 mm isotropic), the equi-volume algorithm can severely amplify discretization and segmentation noise due to its sensitivity to local curvature estimates, and empirical comparisons at this resolution demonstrate that partial voluming effects dominate, rendering the differences between equi-volume and equi-distance layering negligible for functional outcomes [59].

Functional MRI data were first preprocessed by motion correction and slice-timing correction using the FS-FAST pipeline in FreeSurfer. Image distortion due to off-resonance was corrected by estimating field maps from “blip-up” and “blip-down” images [60] (*topup* function in FSL; v. 5.0). Distortion-corrected fMRI was spatially registered to the structural image using boundary-based registration [61] (*bbregister* in FreeSurfer). fMRI data were then sampled onto each depth-specific surface using the *mri_vol2surf* function in FreeSurfer: the pial surface (gray matter/cerebrospinal fluid boundary), the white/gray matter boundary, and nine equally spaced intermediate surfaces between them (n.d.=0.1 to n.d.=0.9, in steps of 0.1). No spatial smoothing was applied between depths. Because gradient-echo EPI signal specificity is degraded by venous signal amplification toward the cortical surface [62], we calculated the average fMRI signal across all surfaces between the pial and white/gray matter boundary and removed it from each depth-specific signal by linear regression.

#### SEEG electrode localization

Electrodes were manually registered to each patient’s MRI using an in-house MATLAB toolbox (fhlin_toolbox, SEEG electrode registration; https://github.com/fahsuanlin/fhlin_toolbox/wiki/SEEG:-register-electrodes-to-MRI-(II)), following our published method [63], applied to the pre- and post-surgery structural MRIs described above. Contacts were assigned to the auditory-cortex ROI for the 15-mm and 5-mm selections based on their registered coordinates relative to the ROI masks used throughout the analysis.

#### SEEG signal extraction

SEEG data were re-referenced to an electrode contact in the white matter, away from any region showing epileptic activity. Frequency-resolved amplitude envelopes were estimated by applying the Morlet wavelet transform (7-cycle width; center frequencies [4:10, 12:2:20, 25:5:60, 70:10:120, 140:20:300] Hz) to the SEEG time series and taking the absolute value of the resulting complex wavelet coefficient at each center frequency, yielding one amplitude-envelope regressor per frequency. Broadband high-frequency activity (HFA), a separate 70–150 Hz band-integrated power measure reflecting non-rhythmic asynchronous neural firing linked to local cortical processing [42,43], was extracted as the squared amplitude of the Hilbert transform of the 70–150 Hz band-pass-filtered SEEG signal. HFA substantially overlaps the wavelet-based high-gamma frequency range (60–300 Hz) and is not an independent regressor from it (see Analysis Models, below).

### Analysis Models

#### SEEG frequency-band models

For each canonical band (alpha: 8–14 Hz; beta: 14–30 Hz; low gamma: 30–60 Hz; high gamma: 60–300 Hz), we fit a separate General Linear Model (GLM) using a single band-specific SEEG regressor — the HRF-convolved amplitude envelope, averaged across the individual wavelet frequencies within that band — together with the nuisance regressors described below. These primary models never combined bands, or SEEG, HFA, and acoustic regressors, within one design matrix; the only exceptions are the nested-model tests and the temporal-derivative control described under Statistical analysis. We refer to this as the canonical-band SEEG model. An analogous single-frequency SEEG model was fit separately at each individual wavelet center frequency.

#### HFA model

HFA envelopes from electrode contacts near the auditory cortex were averaged across contacts and convolved with the same canonical HRF [37], and the resulting single regressor was fit in its own GLM against depth-specific fMRI data, following the same one-regressor-at-a-time design as the SEEG-band models above.

#### Acoustic-envelope model

We extracted the envelope of the presented music using the Hilbert transform (*envelope* function in MATLAB), convolved it with the same canonical HRF, and fit the resulting single regressor in its own GLM against depth-specific fMRI data, following the same one-regressor-at-a-time design. To compare this acoustic-envelope-fMRI coupling profile against the HFA-fMRI coupling profile across depths (**Figure 4B**), we fit the 7-point acoustic-envelope coefficient profile with an exact cubic interpolating spline (MATLAB *fit(…,’cubicspline’)*), evaluated on a fine depth grid, and rescaled it by a single least-squares scalar (fit against the raw 7-point HFA profile) so its amplitude matches the HFA profile it is compared against. The spline curve’s 95% confidence band is a subject-level (n=10) bootstrap that jointly resamples the acoustic-envelope shape and the rescaling factor together. Two limitations apply to this comparison: first, because the scale factor is fit to the same HFA data it is compared against, a close visual match is partly guaranteed by construction and should not be read as independent validation that the acoustic envelope predicts the HFA depth profile; second, even a genuine shape match would not establish that HFA-fMRI coupling reduces to acoustic tracking, because a common, largely frequency- and feature-generic depth-dependent gain (see Results) can make different signals’ depth profiles resemble each other after rescaling, without either being causally reducible to the other.

#### Acoustic-spectrogram model

To test whether SEEG-fMRI coupling and its depth profile could be explained by acoustic tracking alone (see Results), we computed the acoustic spectrogram of the presented music at 84 log-spaced frequency channels (4 Hz–8 kHz), convolved each channel’s envelope with the same canonical HRF, and fit each as its own single-regressor GLM against depth-specific fMRI data, using identical nuisance-regressor and normalization steps as the SEEG-band models above.

#### Nuisance regressors and second-level analysis

In all GLMs, we included six motion parameters and time series from the ventricles and white matter (segmented by FreeSurfer) as nuisance regressors, along with a constant and linear trend. This nuisance structure and two-level design (first-level regression coefficients obtained per participant, followed by a second-level GLM evaluating significance across participants, songs, and repeated presentations) was applied identically to every model described above.

#### Statistical analysis

We particularly focused on the primary (A1) and secondary (A2) auditory cortex defined by the Human Connectome Project data [35]. Because HFA is more spatially local than lower-frequency activity, we analyzed both the primary electrode selection ( contacts within 15 mm of the auditory-cortex centroid) and, for HFA specifically, a more spatially restricted selection (contacts within 5 mm), reporting both where feasible. All statistical tests controlled the false discovery rate (FDR) using the Benjamini-Hochberg (BH) procedure [64]; nested-model comparisons across the four canonical frequency bands instead used the Holm-Bonferroni step-down procedure to control the family-wise error rate [65]. To compare coupling magnitude between regions (e.g., A1 vs. A2) or between conditions on a scale-free basis, each regression t-statistic was converted to a partial correlation (r = sign(t)·√(t²/(t²+df))) and Fisher-z transformed (z = atanh(r)) before averaging or testing across subjects; cross-region and cross-condition magnitude comparisons used these Fisher-z values with a subject-level paired t-test (n=10) rather than a vertex-level test, and used unit-Euclidean-norm-rescaled depth profiles, not raw coefficients, whenever comparing profile shape independent of overall signal scale. Effect sizes for paired comparisons were reported as Cohen’s dz (dz = t/√n, where n is the number of subjects) [66].

#### Nested-model tests

To test whether a regressor carried a unique association with depth-resolved fMRI signals, we compared, for each participant, region, and depth, a full GLM containing the regressor of interest with a reduced GLM omitting it, using an F-test, and assessed the improvement in fit across the 10 participants (F(1,9)), Holm-corrected across the four canonical bands. Two comparisons were made: (i) each canonical band against a reduced model containing the other three bands, and (ii) each canonical band against a reduced model containing the acoustic-feature regressors (broadband envelope plus 8 spectral bands, 50–8,000 Hz). For the temporal-derivative control of the depth-profile shape test, the temporal derivative of each SEEG regressor was added to its GLM.

#### Depth-profile shape test

To test whether two frequency bands differ in the shape of their depth profiles, independent of amplitude and sign, we used an antisymmetric statistic computed for each subject s: T□= (**u□y□**,□)(**v□y□**,□) − (**v□y□**,□)(**u□y□**,□), where **y□**,□ and **y□**,□ are the seven-depth coupling profiles of the two bands, **u** is the band-1 profile averaged over the other nine subjects (unit norm), and **v** is a fixed, unit-norm depth contrast. T□ is the determinant of the 2×2 matrix of projections onto **u** and **v**, and is exactly zero when **y□**,□ = c·**y□**,□ for any scalar c. Writing **v** = (**u□v**)**u** + **v_□_** , the component of **v** along **u** cancels, so T□ depends only on **v_□_** , the part of **v** not shared with the common profile. The contrast **v**, specified in advance, was the expected depth pattern of activity confined to layers III–IV under a published venous-drainage model [38] and the BigBrain depth-to-layer mapping [53]. In A1, **u□v** ≈ 0.97, so **v_□_** carries about 5% of the energy of **v** and alternates in sign across depths. The drainage model only defines **v**; it is not applied to the data, and the validity of the test does not depend on it. Two further contrasts from the same model (layers I–II and V–VI, and its difference from the III–IV pattern) were examined as secondary and were not significant in any region (A1: p=0.95 and p=0.52). Group significance was assessed with a two-sided exact sign-flip permutation test over the 10 subjects (1,024 permutations). Robustness was examined by scaling the drainage model (×0.5–1.5; p≤0.004), by 20 random depth-to-layer mappings (p≤0.010), by leaving out one subject at a time (p=0.004–0.059), and by refitting the GLM with the temporal derivative of each SEEG regressor added, using both the canonical coefficient and the derivative-boosted amplitude [67].

To assess the sensitivity of depth-profile estimates to potential deep-to-superficial venous signal leakage, we additionally performed a deveining sensitivity analysis (our own construction, not a replication of a specific published method). For each depth d (except the deepest, white matter), we regressed that depth’s raw fMRI time series on its immediately deeper neighbor’s raw time series, and applied the correction x ^dev^(t; λ) = x_d_(t) − λ·β ·x (t), where λ _∈_ {0, 0.25, 0.5, 0.75, 1} is the fraction of this empirically fitted relationship treated as venous leakage and removed (λ=0 reproduces the uncorrected pipeline; λ=1 removes the entire linearly predictable contribution from the deeper neighbor). The deveined time series x ^dev^(t; λ) were passed through the same nuisance regressors and SEEG-band GLM described above, and reported as a sensitivity curve across λ rather than a single ‘corrected’ answer, since no physiologically calibrated λ is available for this dataset.

## Acknowledgments

We thank funding from the following institutes to support this research: Canadian Institutes of Health Research grant PJT 178345 and PJT 496433 (FHL), National Sciences and Engineering Research Council grant RGPIN-2020-05927 (FHL), Canada Foundation for Innovation grant 38913 and 41351 (FHL), MITACS IT25405 (FHL).

## Author Contributions

Conceptualization: HYY, WJK, FHL

Methodology: HJL, HYY, KU, WJK, FHL

Investigation: HJL, JA, FHL

Visualization: FHL

Supervision: WJK, FHL

Writing—original draft: HJL, JA, FHL

Writing—review & editing: HJL, JA, HYY, CCC, CCL, WJK, HL, KU, FHL

