## Supplementary material for "Relating layer fMRI signals to acoustics and intracranial neuronal activity in the human auditory cortex in a naturalistic design": supplmental figures and table.

Figures S1 to S2
Tables S1
SI References

### Figures

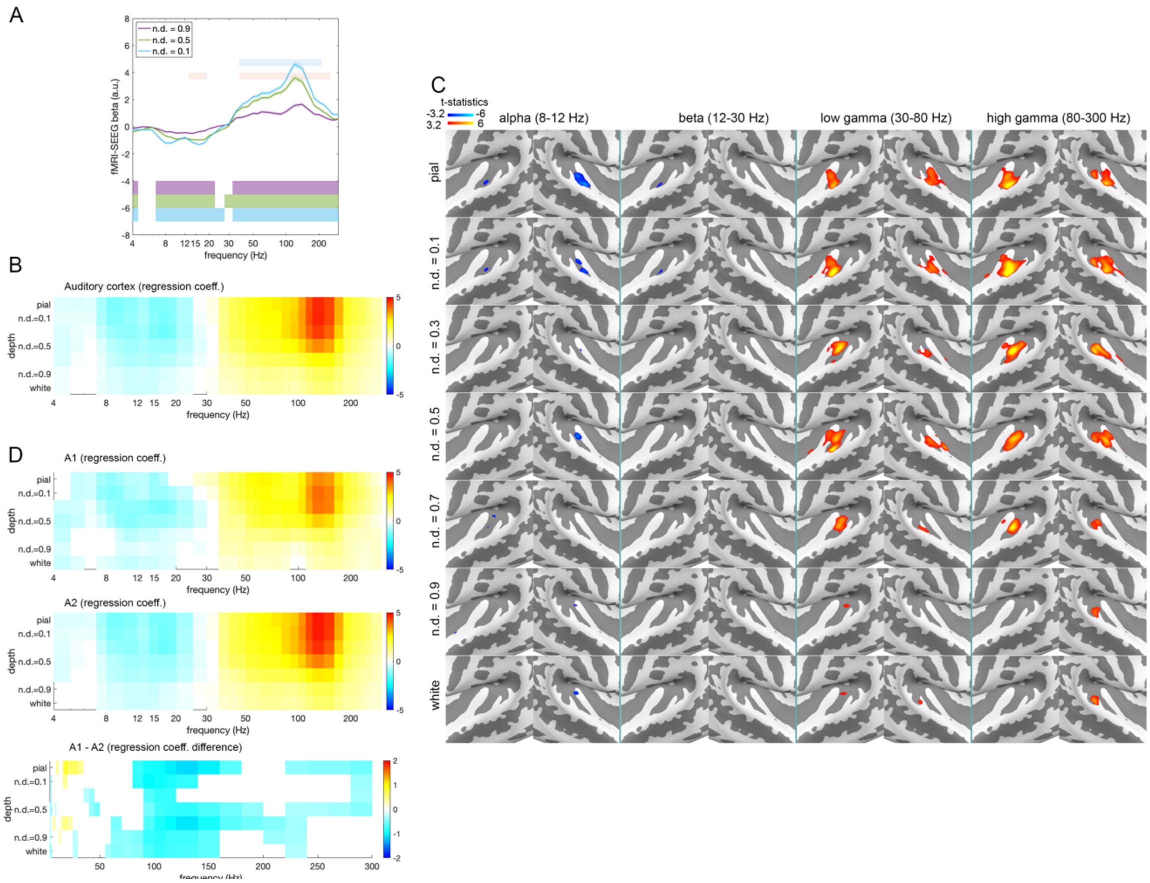


**Fig S1 Correlations between intracranial neural activity and fMRI signals across cortical depths. (A)**: The correlation between fMRI signals at three cortical depths (normalized depth, n.d., of 0.1, 0.5, and 0.9) and neural activity measured by electrodes within 5 mm to the centroid of the auditory cortex. Values are regression coefficients. Significances of regression coefficients (beta's; false discovery rate, FDR, corrected p<0.05 across frequencies) at cortical depths are shown in horizontal bars at the bottom of the panel. The light blue and red horizontal bars at the top of the panel indicate significant differences between the three depths shown in this panel only (n.d.=0.5 vs. n.d.=0.9, and n.d.=0.5 vs. n.d.=0.1, respectively; FDR corrected p<0.05 across frequencies), not a comprehensive depth-pairwise comparison. (**B)**: Regression coefficients between fMRI signals across cortical depths and neural activity. Significant coefficients (FDR corrected p<0.05 across frequencies and cortical depths) are color-coded. (**C)**: Distributions of the significance of correlations between intracranial neural activity and fMRI signals across cortical depths (FDR corrected p<0.05) in alpha, beta, low gamma, and broadband high frequency activity around the auditory cortices in both hemispheres. See **Figure 1** for the enlarged areas. (**D)**: Correlations between neural activity and fMRI signals across cortical depths at primary (A1) and secondary (A2) auditory cortices and their differences. Significant coefficients (FDR corrected $p<0.05$ across frequencies and cortical depths) are color-coded at the vertex level.

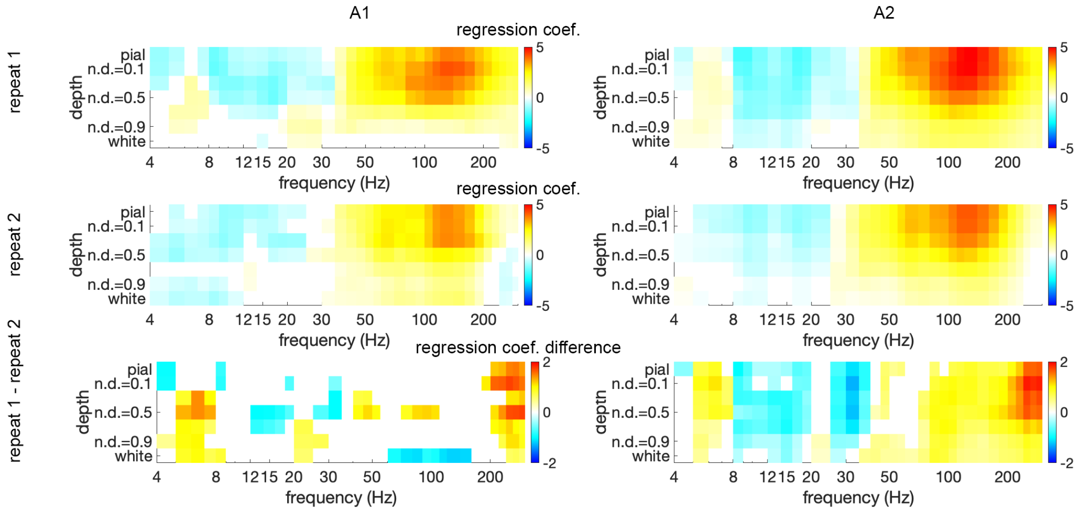


**Fig. S2 Relationships between intracranial neuronal activity and fMRI signals across cortical depths at A1 (left column) and A2 (right column) between repeated listening.** Rows show SEEG-fMRI coupling in the first presentation, the second presentation, and their difference; color-coded cells are significant coefficients (FDR corrected p<0.05 across frequencies and cortical depths) in the first two rows, and significant differences between presentations (vertex-level test, FDR corrected p<0.01) in the bottom row. This comparison tests whether repeated listening differentially affects SEEG-fMRI coupling across cortical depth, frequency band, and region (A1 vs. A2); it does not distinguish neural adaptation from repetition-related changes in neurovascular coupling, and because the SEEG and fMRI measurements come from different participant groups, it cannot establish within-person neural adaptation.

### Tables

**Table S1. MNI coordinates of the implanted electrodes and the distance to the centroid of the auditory cortex in the left (A) and right (B) hemispheres.** Values in Contact column denote the electrode contact from the tip of the implanted electrode. Contacts on the same electrode were separated by 5 mm. The auditory cortex region was defined from an atlas [1].

**A.**

| Patient | Electrode name | Contact | Distance to the center of mass (mm) | X (mm) | Y (mm) | Z (mm) |
|---|---|---|---|---|---|---|
| s050 | sT | 3 | 3.2 | -46.2 | -26.8 | 3.7 |
| s050 | sT | 4 | 4.7 | -51.7 | -25.8 | 3.3 |
| s011 | sT | 3 | 5.0 | -48.8 | -26.1 | -0.1 |
| s011 | sT | 2 | 6.0 | -43.8 | -26.6 | 0.3 |
| s026 | sT | 4 | 6.2 | -48.1 | -29.8 | 2.5 |
| s026 | sT | 5 | 7.7 | -53.0 | -29.0 | 2.6 |
| s050 | sT | 2 | 7.9 | -40.7 | -27.9 | 4.2 |
| s012 | sT | 4 | 8.0 | -47.1 | -31.8 | 5.5 |
| s011 | sT | 4 | 8.1 | -53.8 | -25.7 | -0.5 |
| s026 | sT | 3 | 8.2 | -43.1 | -30.6 | 2.4 |
| s014 | sT | 3 | 9.0 | -46.2 | -32.6 | 2.4 |
| s012 | sT | 5 | 9.1 | -51.7 | -31.9 | 5.3 |
| s012 | sT | 3 | 9.4 | -42.5 | -31.7 | 5.7 |
| s014 | sT | 4 | 9.6 | -51.2 | -32.4 | 2.0 |
| s050 | sT | 5 | 9.8 | -57.2 | -24.7 | 2.9 |
| s011 | sT | 1 | 9.9 | -38.8 | -27.0 | 0.7 |
| s014 | sT | 2 | 11.1 | -41.1 | -32.8 | 2.7 |
| s026 | sT | 6 | 11.4 | -58.0 | -28.2 | 2.8 |
| s012 | sT | 6 | 12.0 | -56.3 | -32.0 | 5.1 |
| s026 | sT | 2 | 12.1 | -38.2 | -31.4 | 2.3 |
| s014 | sT | 5 | 12.4 | -56.3 | -32.2 | 1.7 |
| s012 | sT | 2 | 12.5 | -37.9 | -31.6 | 5.8 |
| s011 | sT | 5 | 12.5 | -58.8 | -25.3 | -0.9 |
| s048 | sE | 4 | 13.1 | -48.0 | -18.7 | 16.2 |
| s050 | sT | 1 | 13.3 | -35.2 | -29.0 | 4.6 |
| s048 | sE | 5 | 13.8 | -53.2 | -18.5 | 15.5 |
| s014 | sY | 3 | 13.8 | -35.2 | -17.7 | 5.6 |
| s048 | sE | 3 | 14.4 | -42.8 | -18.9 | 16.9 |
| s026 | sY | 4 | 14.5 | -38.0 | -13.1 | 6.5 |
| s014 | sT | 1 | 14.7 | -36.0 | -33.0 | 3.1 |
| s014 | sY | 4 | 14.9 | -35.1 | -23.4 | 12.5 |
| s011 | sY | 5 | 14.9 | -33.6 | -21.3 | 9.1 |
| s011 | sY | 4 | 14.9 | -34.2 | -17.0 | 3.9 |

**B.**

| Patient | Electrode name | Contact | Distance to the center of mass (mm) | X (mm) | Y (mm) | Z (mm) |
|---|---|---|---|---|---|---|
| s038 | sT' | 1 | 4.7 | 44.6 | -23.8 | 2.2 |
| s051 | sT' | 4 | 4.8 | 49.7 | -22.5 | 1.0 |
| s038 | sT' | 2 | 4.8 | 48.6 | -20.8 | 1.5 |
| s051 | sT' | 3 | 5.1 | 44.7 | -24.6 | 1.8 |
| s054 | sT' | 4 | 7.4 | 47.4 | -29.6 | 9.7 |
| s038 | sT' | 3 | 8.6 | 52.5 | -17.9 | 0.8 |
| s051 | sT' | 5 | 9.0 | 54.7 | -20.5 | 0.2 |
| s051 | sT' | 2 | 9.3 | 39.7 | -26.6 | 2.6 |
| s047 | sT' | 2 | 10.2 | 47.1 | -32.9 | 1.0 |
| s054 | sU' | 3 | 10.8 | 48.9 | -15.9 | -2.1 |
| s036 | sT' | 3 | 10.9 | 46.4 | -32.3 | -1.2 |
| s039 | sY | 6 | 11.0 | 37.6 | -24.3 | 2.0 |
| s047 | sT' | 3 | 11.2 | 52.5 | -32.8 | 0.6 |
| s036 | sT' | 4 | 11.3 | 51.3 | -32.1 | -1.5 |
| s047 | sT' | 1 | 11.8 | 41.8 | -33.0 | 1.5 |
| s054 | sT' | 6 | 11.9 | 57.7 | -29.8 | 8.5 |
| s039 | sY | 7 | 11.9 | 37.2 | -28.2 | 7.4 |
| s036 | sY' | 6 | 12.2 | 36.6 | -19.9 | 6.8 |
| s054 | sU' | 4 | 12.5 | 54.1 | -16.5 | -2.9 |
| s036 | sT' | 2 | 12.5 | 41.5 | -32.5 | -0.9 |
| s038 | sT' | 4 | 13.2 | 56.5 | -14.9 | 0.1 |
| s036 | sT' | 5 | 13.6 | 56.2 | -31.9 | -1.8 |
| s031 | sT' | 2 | 13.6 | 43.6 | -19.2 | -6.7 |
| s039 | sY | 5 | 13.8 | 38.0 | -20.4 | -3.4 |
| s036 | sY' | 5 | 13.8 | 37.1 | -15.9 | 2.5 |
| s051 | sT' | 6 | 14.2 | 59.8 | -18.4 | -0.7 |
| s047 | sT' | 4 | 14.3 | 57.9 | -32.7 | 0.1 |
| s051 | sT' | 1 | 14.5 | 34.6 | -28.7 | 3.4 |
| s031 | sT' | 4 | 15.0 | 55.4 | -17.0 | -5.9 |
